# Zebrafish and cellular models of *SELENON*-Related Myopathy exhibit novel embryonic and metabolic phenotypes

**DOI:** 10.1101/2024.02.26.581979

**Authors:** Pamela Barraza-Flores, Behzad Moghadaszadeh, Won Lee, Biju Isaac, Liang Sun, Emily C. Troiano, Shira Rockowitz, Piotr Sliz, Alan H. Beggs

## Abstract

*SELENON*-Related Myopathy (*SELENON*-RM) is a rare congenital myopathy caused by mutations of the *SELENON* gene characterized by axial muscle weakness and progressive respiratory insufficiency. Muscle histopathology commonly includes multiminicores or a dystrophic pattern but is often non-specific. The *SELENON* gene encodes selenoprotein N (SelN), a selenocysteine-containing redox enzyme located in the endo/sarcoplasmic reticulum membrane where it colocalizes with mitochondria-associated membranes. However, the molecular mechanism(s) by which SelN deficiency causes *SELENON*-RM are undetermined. A hurdle is the lack of cellular and animal models that show assayable phenotypes. Here we report deep-phenotyping of SelN-deficient zebrafish and muscle cells. SelN-deficient zebrafish exhibit changes in embryonic muscle function and swimming activity in larvae. Analysis of single cell RNAseq data in a zebrafish embryo-atlas revealed coexpression between *selenon* and genes involved in glutathione redox pathway. SelN-deficient zebrafish and mouse myoblasts exhibit changes in glutathione and redox homeostasis, suggesting a direct relationship with SelN function. We report changes in metabolic function abnormalities in SelN-null myotubes when compared to WT. These results suggest that SelN has functional roles during zebrafish early development and myoblast metabolism.

## INTRODUCTION

*SELENON*-Related Myopathy (*SELENON*-RM) is a rare congenital myopathy caused by mutations of the *SELENON* gene (Ferreiro et al., 2002; Moghadaszadeh et al., 2001). It has an estimated incidence of 1 in 200,000 births (Bouman et al., 2021; Witting et al., 2017). However, based on population estimates of carrier frequencies the incidence may be significantly higher when considering undiagnosed cases. *SELENON*-RM has a clinical onset mostly before two years of age but may also present as late as early to mid-adulthood (de Visser, 2020; Silwal et al., 2020). It is characterized by the development of early-onset rigidity of the spine, axial muscle weakness, scoliosis, and respiratory insufficiency requiring nocturnal ventilation; however, the great majority of patients remain ambulant throughout their lives. There is currently no cure or specific treatment for *SELENON*-RM patients. Patients’ muscle biopsies show heterogenous histopathological changes that have led to the classification of this disease as rigid spine muscular dystrophy (RSMD) (Moghadaszadeh et al., 2001), multiminicore disease (Ferreiro et al., 2002), congenital fiber type disproportion (Clarke et al., 2006), and Desmin-related myopathy with Mallory-like body inclusions (Ferreiro et al., 2004; Schara et al., 2008). However, these changes are not specific and are not universally found in patients. The most common histopathological findings are multiminicore lesions, which reflect areas of mitochondrial loss (Villar-Quiles et al., 2020). Additionally, natural history studies in *SELENON*-RM patients report insulin-resistance (Varone et al., 2019) as well as altered body mass index, both of which correlate with disease severity (Filipe et al., 2021; Scoto et al., 2011).

The *SELENON* gene encodes selenoprotein-N (SelN), which is a member of the selenoprotein family. Selenoproteins are a group of selenocysteine-containing proteins involved primarily in oxidoreduction reactions. Their selenocysteine residues are involved in enhancing redox catalytic activity (Labunskyy et al., 2014). SelN is an endo/sarcoplasmic reticulum (ER/SR)-resident glycoprotein with a transmembrane domain (Petit et al., 2003a), thiol reductase catalytic activity, and a calcium binding EF-hand domain (Chernorudskiy et al., 2020). SelN has been proposed as an important ER/SR redox protein that plays an essential role in cell protection against oxidative stress and modulation of SERCA channel activity (Arbogast et al., 2009; Pozzer et al., 2019; Zito and Ferreiro, 2021). Previous studies suggest that inducing ER stress in SelN-deficient myoblasts can contribute to increased pathological phenotypes (Pozzer et al., 2019; Varone et al., 2019). SERCA channels are redox partners of SelN, and lack of SelN leads to SERCA channel oxidation and inactivation, resulting in reduced calcium uptake to the ER/SR. While this mechanism likely contributes to delayed myofiber relaxation(Pozzer et al., 2019), it is probably not the only pathophysiological mechanism leading to weakness in *SELENON*-RM patients. It has been shown that SelN also colocalizes with Mitochondria Associated Membranes (MAMs) that tether the mitochondria to the ER (Filipe et al., 2021). MAMs allow for Ca^2+^ influx from the ER into the mitochondria, phospholipid synthesis and transport, regulation of mitochondrial dynamics, among other vital cell functions (Hemel et al., 2021; Nieblas et al., 2022). Based on these findings, we hypothesize that mitochondria dysregulation may play a vital role in the disease mechanism of *SELENON*-RM.

*SELENON* is expressed broadly in mature tissues throughout the body, but its highest levels of expression are during early developmental stages (Petit et al., 2003b). *In situ* hybridization studies have shown that *Selenon* transcripts in mouse and zebrafish embryos are located in developing notochord and somite regions (Castets et al., 2009; Deniziak et al., 2007; Petit et al., 2003b; Thisse et al., 2003; Wagner et al., 2018), suggesting an important role during early muscle development. Even though there is an identified pattern of expression during embryogenesis and some molecular functions for SelN, one of the main obstacles for the study of *SELENON*-RM has been an inability to demonstrate a robust disease phenotype in animal and cellular models. Previously, our group developed a *Selenon* null mouse model for the study of this disease (Moghadaszadeh et al., 2013). We found changes in lung development and mild core lesions in the muscle could be observed only when the mice were subjected to oxidative stress via vitamin E deficiency and forced endurance running. Other groups have shown a fatigue-induced phenotype in ex vivo mechanical studies of a similar *Selenon* null mouse model (Rederstorff et al., 2011), however, clinical signs of myopathy were not readily apparent.

Although cellular models of *SELENON*-RM have been used to describe molecular mechanisms of SelN (Arbogast et al., 2009; Chernorudskiy et al., 2020; Filipe et al., 2021; Pozzer et al., 2019), robust functional assays that might become the basis for small molecule screens have been difficult to develop. Several morpholino-based transient models of SelN knockdown in zebrafish have been reported (Deniziak et al., 2007; Jurynec et al., 2008), but it is unclear what aspects of the phenotypes were due to SelN deficiency and what might be off-target consequences of the morphoplinos. No stable zebrafish models of germline *selenon* knock-out have been reported to date. In this project, we aim to deep-phenotype new zebrafish and cellular knock-out models for the study of *SELENON*-RM while elucidating SelN functional roles in muscle metabolism.

## RESULTS

### Zebrafish models of *SELENON*-RM show null or reduced expression of SelN

Using CRISPR-Cas9 technology we established three independent zebrafish lines with different mutations in exon-2 of the *selenon* gene (NM_001004294.4). Two mutations, selenon:c.197-200del (ZFIN zebrafish line *selenon*^cl502^) and c.199-203del (*selenon*^cl503^), remove four and five base pairs respectively, leading to frameshifts p.L67RfsX57 and p.L67GfsX16 predicted to result in early truncation of the protein, representing presumptive null mutations of SelN (*selenon-*KO). The third is a 3 base pair deletion, c.197-199del (*selenon*^cl504^), resulting leading to in-frame loss of one amino acid, p.G66VdelL67 **(Fig. 1A).** Three-D modeling of zebrafish wild type (WT) and p.G66VdelL67 SelN using SWISS-MODEL (Guex et al., 2009; Waterhouse et al., 2018) predicts disruption of an α-helix in the N-terminal domain of SelN **(Supplemental Fig. 1)**.

**Figure 1.**
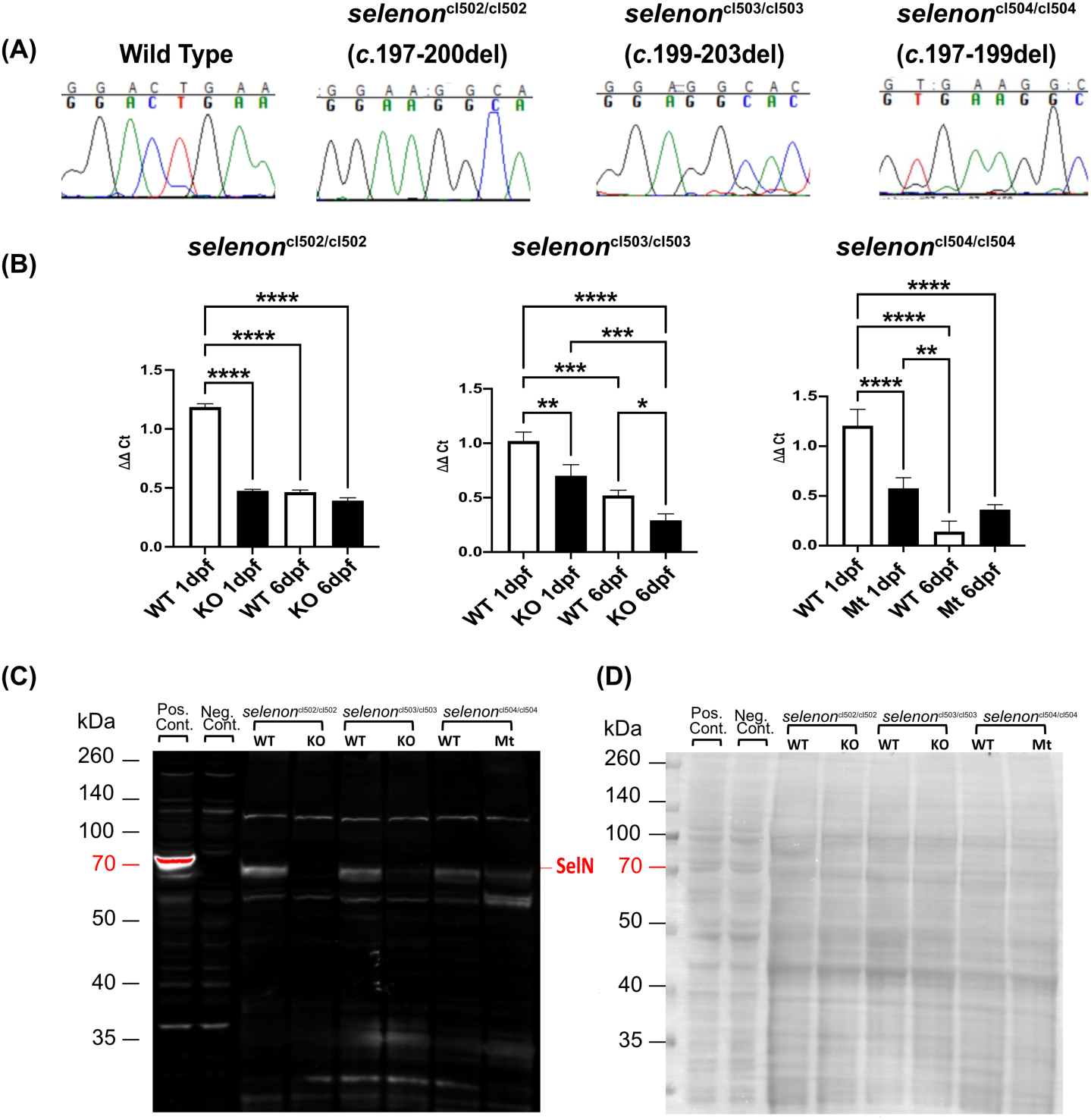
Zebrafish *selenon* mutants show absent or partial expression of Selenoprotein-N. **(A)** Sanger sequencing chromatograms show analysis of the *selenon* gene exon 2 for wild type (WT), *selenon*^cl502^, *selenon*^cl503^, and *selenon*^cl504^ homozygotes using genomic DNA from zebrafish tail clips. **(B)** Real Time qPCR analyses show *selenon* transcript levels at 1dpf and 6dpf in zebrafish knock outs (KO’s) and mutants (Mt): *selenon*^cl502^, *selenon*^cl503^, and *selenon*^cl504^with their correspondent WT. (N=5 per group) “*” = p<0.05, “**” = p<0.01, “***” = p<0.001, “****” = p<0.0001. **(C)** Western Blot analysis shows SelN expression at the predicted size of ∼65kDa in positive control (SelN-transfected HEK cells) but not in negative control (WT HEK cells). Protein lysates from 2dpf zebrafish *selenon* mutants and their corresponding WT controls show SelN expression in all WT fish, no expression in *selenon*^cl502^ and *selenon*^cl503^, and reduced expression in *selenon*^cl504^ mutant line. (N=30 per group) (D) Ponceau staining in western blot used to demonstrate equal protein loading throughout the blot.

This suggests that the deletion of Leucine 67 affects the folding of SelN and interferes with its normal function. To assess consequences of these mutations we measured relative *selenon* transcript levels in wild type (WT) and homozygous mutants of the three zebrafish lines at 1- and 6-day post-fertilization (dpf) **(Fig. 1B).** The highest levels of *selenon* transcript were found in WT embryos at 1dpf, while same age-mutant embryos showed approximately half the expression. By 6dpf, *selenon* expression in all lines dropped by half or more compared to 1dpf WT embryos. Lastly, 6dpf *selenon-*KO lines showed the same or lower transcript levels when compared to WT, while 6dpf *selenon*^cl504/cl504^ showed a trend towards increased transcript level when compared to WT. Next, we analyzed Selenoprotein-N (SelN) protein content in 2dpf zebrafish **(Fig. 1C, D).** SelN expression was undetectable in the two frameshifted and presumptive null lines (*selenon*^cl502/cl502^ and *selenon*^cl503/cl503)^ and apparently reduced by about 55% in *selenon*^cl504/cl504^ fish with a single amino acid deletion.

### Zebrafish models of *SELENON*-RM exhibit reduced spontaneous contraction during embryogenesis

During zebrafish somitogenesis embryos begin to spontaneously contract their trunk and tail from 17 hours post-fertilization (hpf) to approximately 26hpf (Drapeau et al., 2002; Saint-Amant and Drapeau, 1998). The frequency of these muscle contractions is dependent on motor neuron activation and this process is essential for proper axial skeletal muscle formation (Buss and Drapeau, 2001; Liu and Westerfield, 1988). Lesions in the hindbrain do not inhibit these contractions indicating that these events result from activation of the spinal neurocircuitry with no input from the brain (Saint-Amant and Drapeau, 1998). Additionally, it has been shown that muscle contraction at this stage is dependent only on slow twitch fibers (mitochondrial positive fibers) with no input from fast twitch fibers (Naganawa and Hirata, 2011). Zebrafish embryo transparency allowed us to directly observe and quantify spontaneous contractions as a mean to study embryonic muscle function. Fish embryos at 24hpf were videotaped for 3 min and their spontaneous movements were later analyzed computationally. We quantified mean durations of contraction, percentage of total active time, and total numbers of contractions. *Selenon* mutants exhibited significantly decreased mean duration of contractions across all three lines (**Fig. 2A)**. The percentage of active time was significantly lower in the two *selenon* KO lines when compared to WT; while *selenon*^cl504/cl504^ embryos showed a trend towards decrease when compared to WT. The absolute number of contractions, on the other hand, showed no changes in *selenon*^cl502/cl502^ but was significantly reduced in *selenon*^cl503/cl503^and significantly increased in *selenon*^cl504/cl504^. These results indicate that the mean duration of contraction was the most consistent difference during spontaneous contractions in *selenon* mutated zebrafish embryos. To understand whether these changes in contraction were due to delayed early zebrafish growth, we quantified hatching activity in these three lines **(Fig. 2B).** To do so, we generated embryos from matings of *selenon* heterozygous fish and observed the hatching activity from 3 to 5dpf. Hatching time courses for subsequently genotyped fry of all three lines demonstrate that there are no differences between WT, heterozygous (HET), and *selenon-*KO or *selenon*^cl504/cl504^ fish. These data indicate that SelN-deficiency does not lead to hatching abnormalities or delay in zebrafish.

**Figure 2.**
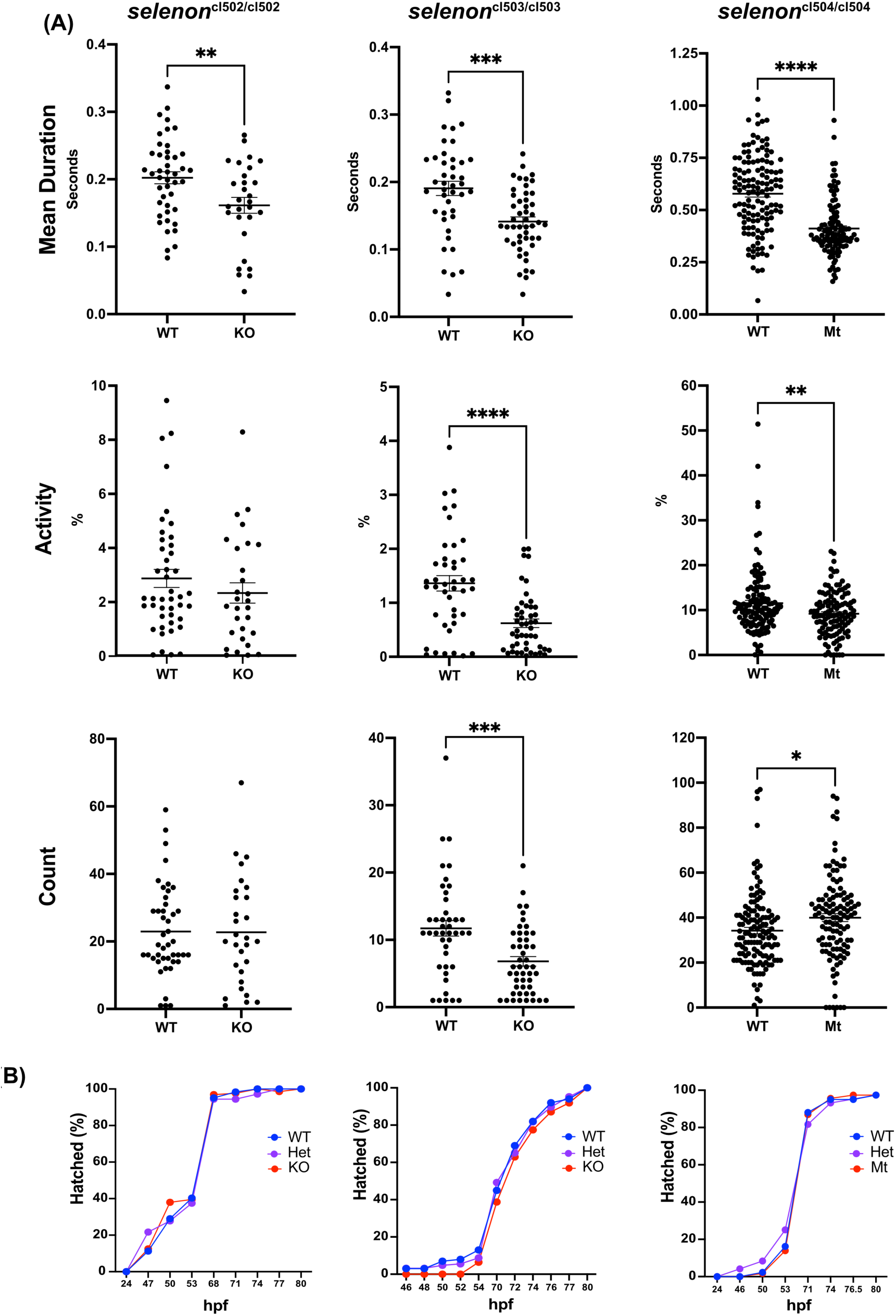
Selenoprotein-N deficient zebrafish embryos present with impaired spontaneous contractions. Spontanous tail and trunk contractions of zebrafish embryos were recorded and quantified at 24hpf in SelN homozygous KO *selenon*^cl502^ (N=45 and 29), *selenon*^cl503^ (N=43,48), and homozygous mutant (Mt) *selenon*^cl504^ (N=131,116) and their correspodent WT controls. **(A)** Mean duration of spontaneous contractions, percent time of contraction activity, and total number of contractions are reported in each line. “*” = p<0.05, “**” = p<0.01, “***” = p<0.001, “****” = p<0.0001. **(B)** WT, heterozygous and homozygous embryos were observed for hatching activity and recorded up until 80hpf (N = 41-100 WT, 92-134 heterozygous, and 56-93 homozygous mutant embryos per line).

### Selenoprotein N-deficient zebrafish larvae show changes in swim activity

To assess motor activity during larval stages of development we tested our SelN*-*deficient zebrafish lines using a swim activity monitor. 6dpf larvae were evaluated using a protocol that included vibrations and light-and-dark cycles. As an example, we show the swimming activity of *selenon*^cl502/cl502^ and WT zebrafish throughout the protocol **(Fig. 3A)**. Changes in light and vibration induced continuous swim activity in both WT and KO, but overall, the WT zebrafish swam more than the KO. Therefore, we focused on quantifying the total activity of the different fish lines. *Selenon* KO lines showed significant decreases in total swim activity when compared to WT, while *selenon*^cl504/cl504^ showed a trend towards decreased swim activity (**Fig. 3B)**.

**Figure 3.**
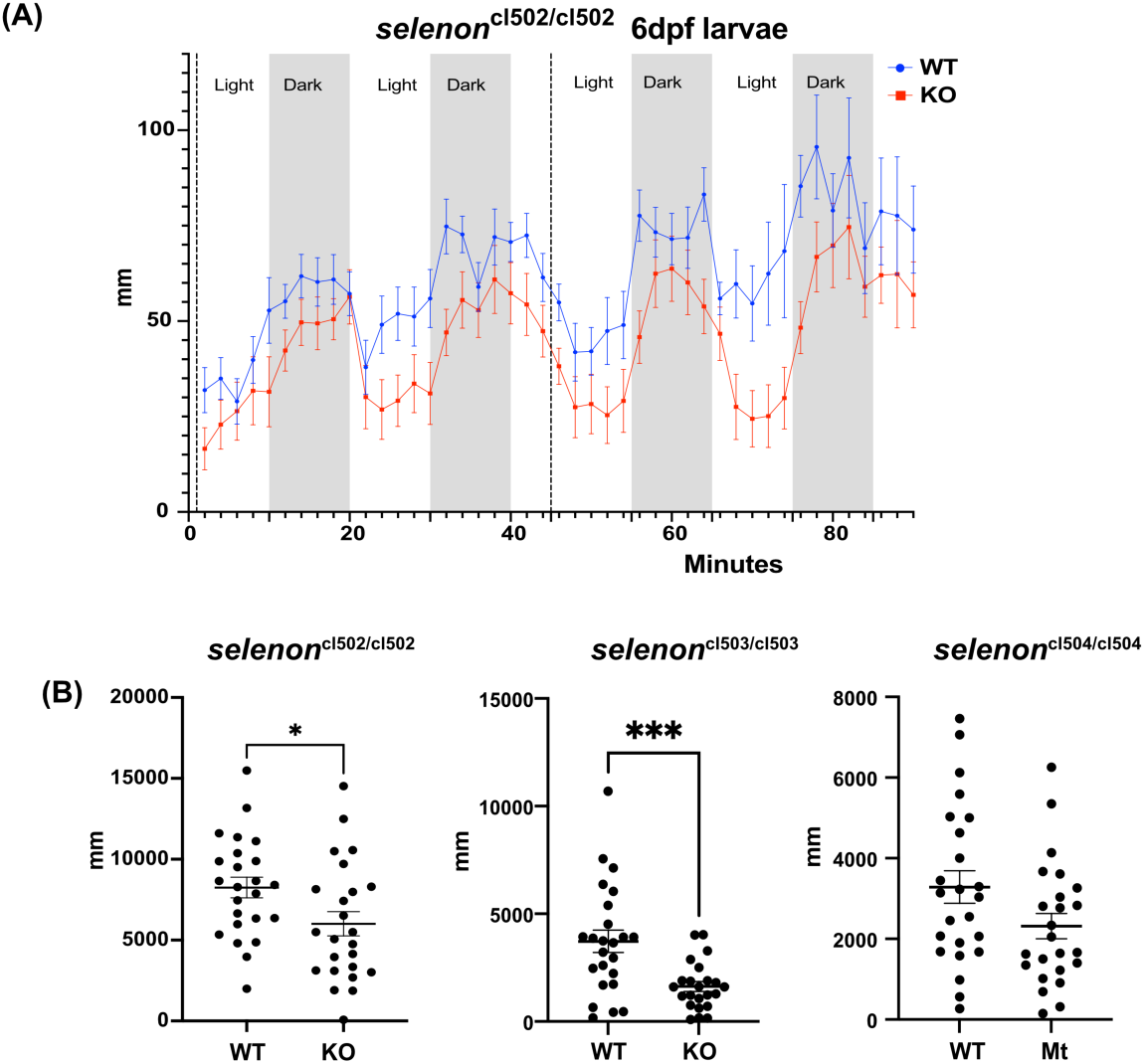
*Selenon-*deficient zebrafish larvae show decreased activity. **(A)** 6dpf WT (blue) and homozygous *selenon-*KO *selenon^cl502/^*^cl502^(red) zebrafish larvae were tested for swim activity using an activity monitor and a 90-minute protocol that included vibration (dotted lines) and cycles of alternating light (non-shaded areas) and dark (shaded areas) periods. **(B)** Quantification of activity assays of WT and homozygous *selenon*-mutant lines shows decreased total distance swum in SelN*-*deficient zebrafish larvae when compared to their correspondending WT controls (N=24). “*” = p<0.05, “***” = p<0.001.

To determine whether these changes in activity were due to ultrastructural modifications in the zebrafish larvae skeletal muscle, we imaged 6dpf *selenon-*mutants and their corresponding WT using transmission electron microscopy (TEM). We show representative images of WT and KO *selenon*^cl502^ skeletal muscle myofibrils **(Supplemental Fig.2)**. We did not identify changes in contractile structures. However, we quantified intermyofibrillar mitochondria area and observed a significant increase in SelN-KO fish lines when compared to WT and a decrease in *selenon*^cl504/cl504^when compared to WT.

### Analysis of gene expression in a zebrafish single cell atlas identifies correlation between *selenon* gene expression and ER oxidoreductase activity in skeletal muscle precursor cells

To understand patterns of *selenon* expression during zebrafish early development we consulted single cell RNAseq atlases by Wagner *et al*. 2017 and Farnsworth *et al.,* 2020 (Farnsworth et al., 2020; Wagner et al., 2018). The Wagner publication sequenced cell transcripts from 1 to 24hpf, while the Farnsworth publication sequenced cells from 1dpf to 5dpf. *Selenon* patterns of expression in both atlases revealed peak expression at 24hpf in both notochord and tailbud presomitic mesoderm cells (PSM), which are precursors of skeletal muscle. This is consistent with previous *in vitro* hybridization studies in mouse and zebrafish embryos (Castets et al., 2009; Petit et al., 2003b; Thisse et al., 2003; Wagner et al., 2018). Based on this, we focused our attention on Wagner’s atlas tailbud PSM cell clusters. In this atlas, over 90,000 cells’ transcriptomes were sequenced throughout zebrafish embryonic development. These transcriptomes were then clustered and constructed in a tree-like diagram that branches into the different trajectories of zebrafish cell differentiation **(Fig. 4A).** The analysis reveals incremental expression increases in tailbud PSM cells at 6, 8, 10, 14, 18, and 24hpf cell clusters. The transcriptome data from the highlighted PSM clusters was used to identify correlations between expression of *selenon* and other genes. Weighted correlation networks were generated using the WGCNA method with the transcriptome data (see Methods) and two modules, ‘antiquewhite1’ (4516 genes) and ‘firebrick2’ (240 genes) were selected based on target selenon gene having significant correlation with p-values < 0.05 **(Supplemental Files 1 and 2, correspondingly)**. Using Cytoscape, 3339 genes from ‘antiquewhite1’ and 168 from ‘firebrick2’ were visualized as *selenon* positively associated gene networks. All co-expressed genes from significant modules ‘antiquewhite1’ and ‘firebrick2’ were subjected to pathway and gene ontology (GO) enrichment analysis. Following pathway enrichment, GO analysis revealed pathways involved in molecular functions, cellular components, and biological processes. Due to its relevance to SelN’s oxidoreductase activity, we focused on the ‘oxidoreductase activity acting on peroxide as an acceptor’ GO pathway in ‘antiquewhite1’ module. **Supplemental table 1** lists the genes included in this group organized by weight of correlation to *selenon*. To further explore the genes listed in **Supplemental Table 1**, we utilized Ingenuity Pathway Analysis (IPA) to identify potential pathways involved with SelN function. We show IPA identified pathways upregulated with SelN organized by statistical significance **(Fig.4B)**. “Glutathione Redox Reactions I” and “Apelin Adipocyte Signaling Pathway” head the list with positive z-scores indicating gene upregulation. We focused on “Glutathione Redox Reactions I” which involves 4 redox genes: *gpx7, gpx8, gpx4a and gpx1a* **(Fig. 4C)**.

**Figure 4.**
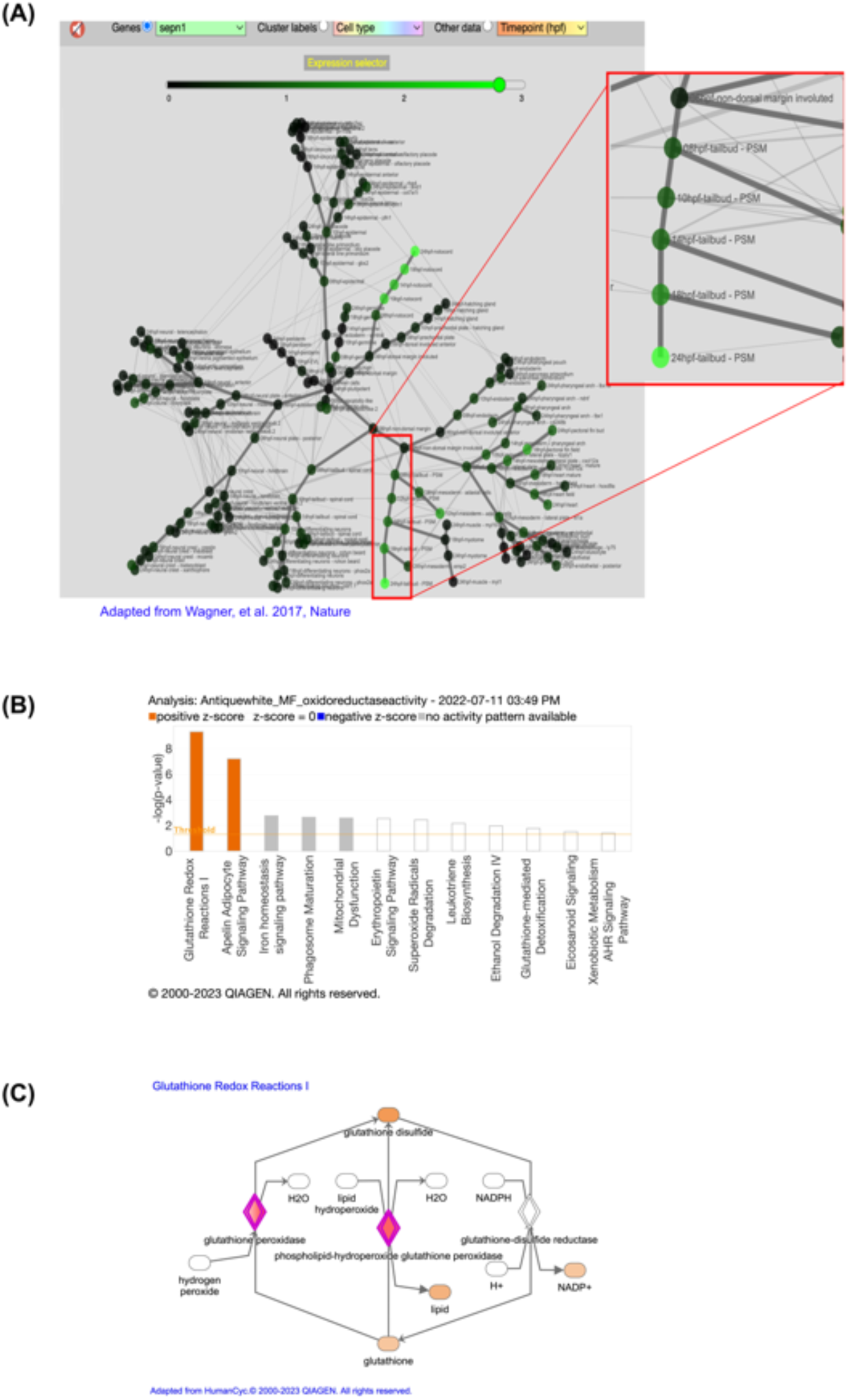
Analysis of embryonic single cell RNAseq zebrafish atlas reveals correlation between *selenon* expression and glutathione redox genes in tailbud presomitic mesoderm (PSM). **(A)** Tree-cell cluster diagram from Wagner et al. 2017, shows *selenon* expression (green) throughout early embryogenesis. Tailbud PSM cell clusters show incremental expression of *selenon* from 6 to 24hpf (red square). **(B)** Pathway analysis of genes coexpressed with *selenon* identified in “oxidoreductase activity” gene ontology reveal overexpression of glutathione redox genes as well as apelin adipocyte signaling pathway. **(C)** Glutathione redox pathway diagram shows the four highly correlated genes with *selenon* (purple): *gpx7, gpx8, gpx4a,* and *gpx19*.

### Glutathione redox homeostasis is altered in SelN deficient cell and zebrafish models

Based on our observation using the single cell RNAseq zebrafish atlas, we hypothesized that co-expression of *selenon* with components of the glutathione redox pathway may reflect functional relationships. To test this, we induced *Selenon* transcript knock down in C2C12 mouse myoblasts using shRNA. *Selenon* transcript levels in three C2C12 knock down (KD) cell lines were reduced to ∼20% in line KD1, ∼10% in KD8 and ∼60% in KD3 relative to parallel control lines **(Supplemental Fig. 3)**. By western blotting, SelN protein expression was overall reduced in KD1 and KD8 cell lines, relative to controls.

Next, we assayed our SelN-KD cell lines and SELENON-RM zebrafish models for glutathione (GSH) redox states. In our cell lines, we quantified the ratio between oxidized (GSSG) and reduced (GSH_free_) forms of GSH. This ratio allowed us to identify global changes in glutathione redox state. Results show that the glutathione ratio (GSH_free_/GSSG) was significantly downregulated in both KD1 and KD8 and maintained the same trend towards decreased ratio in KD3 when compared to control (**Fig. 5A)**. This suggests that GSH redox ratio is sensitive to downregulation of the SelN protein in C2C12 myoblasts.

**Figure 5.**
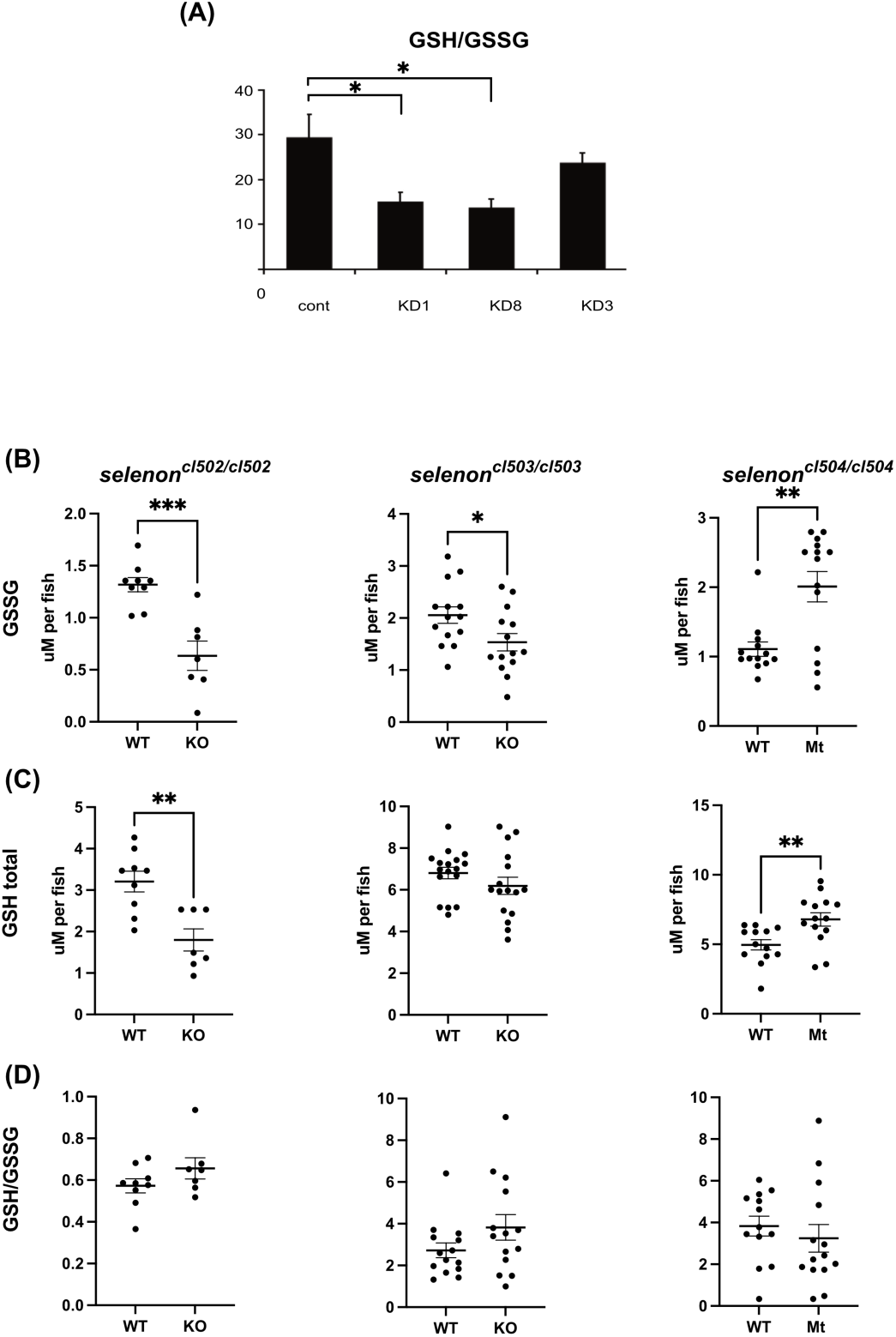
Glutathione homeostasis is altered in cell an zebrafish models of SELENON-Related Myopathy. **(A)** *Selenon* Knock Down (KD) cell lines show decreased gluathione ratio (GSH/GSSG) when compared to WT control. **(B)** GSSG, **(C)** total GSH, and **(D)** GSH/GSSG ratio were measured in 6dpf *selenon*^cl502^ (N=9 and 7), *selenon*^cl503^ (N=14), and *selenon*^cl504^ (N=13 and 14) zebrafish lines. Results show altered glutathione homeostasis. “*” = p<0.05, “**” = p<0.01, “***” = p<0.001.

Additionally, we tested GSH redox states in our zebrafish models of SELENON-RM at 6dpf. Due to ease of normalization with fish, we were able to report the total amount of GSH (GSH_total_), GSSG, as well as the GSH_free_/GSSG ratio. We show that levels of GSSG were significantly reduced in the *selenon*-KO fish when compared to their corresponding WT fish **(Fig. 5B)**. However, *the selenon^cl504/cl504^* fish showed significantly increased levels of GSSG compared to WT. We also show a significant reduction in GSH_total_ in the *selenon^cl502/cl502^* fish line while the *selenon^cl503/cl503^*showed a trend towards decreased levels **(Fig. 5C)**. Additionally, the *selenon^cl504/cl504^* fish showed a significant increase in GSH_total_ when compared to WT. Finally, we show the GSH_free_/GSSG ratios in each fish line **(Fig. 5D)**. Results show no significant differences between any SelN-deficient fish line and their matched WT controls. However, a trend towards increased ratio was observed in the *selenon*-KO lines while the *selenon^cl504/cl504^*fish showed a trend towards decreased ratio. These data suggest that zebrafish GSH redox state at 6dpf is sensitive to levels of SelN protein.

### SelN deficient myoblasts exhibit increased oxidative stress

To determine whether SelN downregulation induces oxidative stress, we quantified the presence of reactive oxygen species (ROS) in SelN-KD and control C2C12 cells. To do so we used CM-H2DCFDA general oxidative stress indicator in our cell lines and flow cytometry. Results show a shift towards increased ROS in both the KD1 and KD8 when compared to control cells **(Fig. 6A)**. We also show quantification of the mean fluorescence per group which shows a significant increase in KD1 and KD8 when compared to control cells **(Fig. 6B)**. To determine whether the higher levels of ROS in SelN-deficient cells are associated with activation of oxidative damage or modification, we quantified levels of carbonylation and nitrosylation in each line. Both SelN-KD lines showed a significant increases in levels of carbonylated proteins relative to controls **(Fig. 6C)**. Similarly, we show an increase in nitrosylated proteins in SelN-KD cell lines when compared to control cells **(Fig. 6D)**. These data indicate that SelN-deficiency induces high levels of oxidative stress resulting in irreversible protein oxidation.

**Figure 6.**
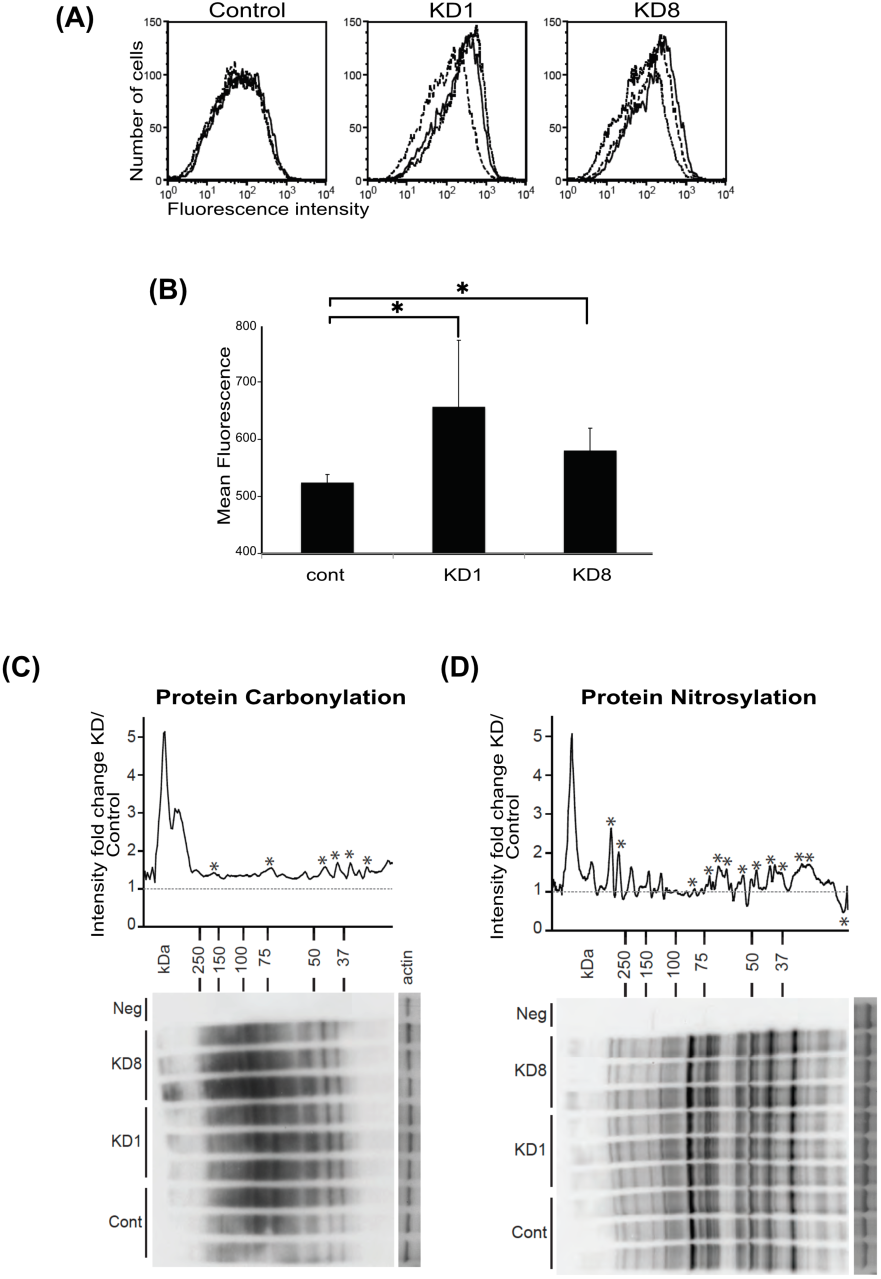
*Selenon* knock down cells present increased ROS levels when compared to control. **(A)** Flow cytometry assay shows fluorescently labeled ROS levels in *selenon* knock down (KD1 and KD8) and control cells. **(B)** Quantification of fluorescence reveals significant increase of ROS levels in *selenon* KD cell lines when compared to control. **(C)** Immunoblotting of protein carbonylation and **(D)** nytrosylation demonstrate increased levels in *selenon* KD when compared to control cells.

### Selenoprotein-N null myotubes show impaired metabolism

Given the correlation between SelN deficiency and mitochondrial abnormalities, we hypothesized that *Selenon*-KO myoblasts may present metabolic impairment. To test this idea, using CRISPR-Cas9 technology we independently created two new C2C12 myoblast lines with homozygous *Selenon* null mutations in exon 3 and exon 5. Additionally, we isolated primary quadriceps myoblast cell lines from our mouse model of *SELENON*-RM, which has a homozygous null mutation in exon 8 (Moghadaszadeh et al., 2013). Each line was tested with a matched WT cell line for comparison. Immunoblot analysis of SelN protein confirmed its absence in both C2C12 cell lines as well as in our mouse Selenon-KO quadriceps primary cells **(Fig. 7A)**.

**Figure 7.**
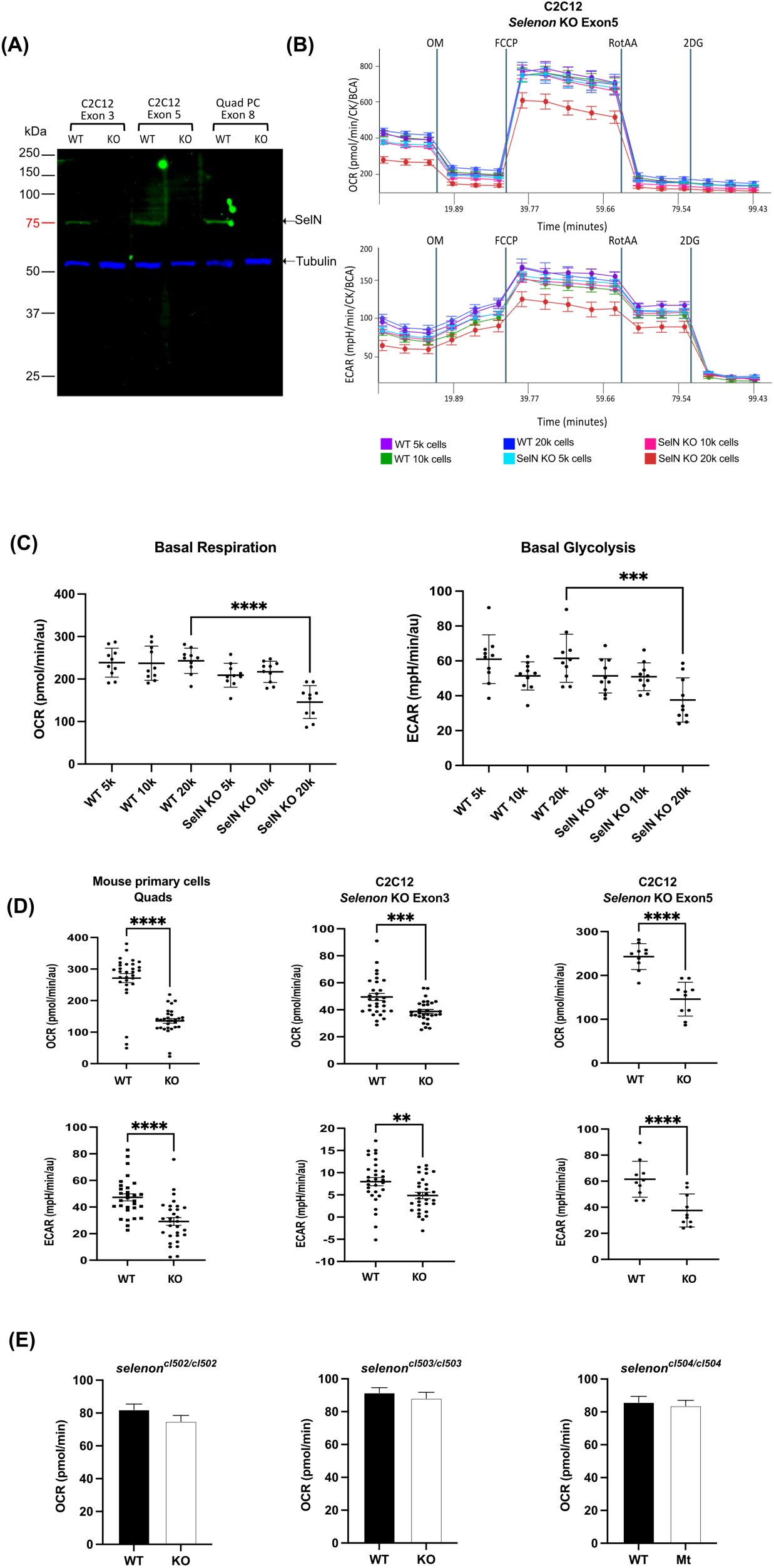
Selenoprotein-N null myoblasts show impairment in metabolism after differentiation at high cell seeding confluency. **(A)** Immunoblot shows absence of ∼65kDa SelN in three *selenon*-null myoblast lines: C2C12 Exon 3, C2C12 Exon 5, and mouse quadriceps primary cells (Quad PC). α-Tubulin was used as a loading control and is present at ∼52kDa in all lines. **(B)** Results from seahorse cell respirometer show changes in Oxygen Consumption Rate (OCR) and Extra Cellular Acidification Rate (ECAR) parameters at different cell densities of *selenon*-null myoblast lines. **(C)** Quantification of C2C12 *selenon*-null exon 5 line’s basal levels of OCR and ECAR show differences in metabolism when compared to wild type at 20,000 cell density (N=10). **(D)** Quantification of basal OCR and ECAR in two C2C12 and one mouse quadricep primary *selenon-*null cell lines shows impaired metabolism in KO myoblasts when compared to WT (N=30). **(E)** Quantification of basal OCR in three 1dpf *selenon*-mutant zebrafish embryos show no differences in metabolism in mutants when compared to WT (N=48). “**” = p<0.01, “***” = p<0.001, “****” = p<0.0001.

To test for changes in metabolism in our cell lines, we used the Seahorse cell respirometer to measure Oxygen Consumption Rate (OCR) and Extra Cellular Acidification Rate (ECAR). We used increasing concentrations of exon 5 *Selenon-*KO and WT myoblasts. Cells were then differentiated into myotubes and measured for OCR and ECAR. After completion, we measured creatine kinase activity and protein concentration in each well for normalization of differentiation and cell number respectively. Using OCR values, we calculated basal levels of cellular respiration, ATPase activity, maximal respiration capacity, and non-mitochondrial oxygen consumption. At the same time, with ECAR levels we calculated basal glycolysis, maximal glycolysis levels, and non-glycolytic acidification (data not shown). Results show an overall decrease in all OCR and ECAR parameters in the *Selenon-*KO myotubes when seeded at 20,000 cells per well and differentiated for five days **(Fig. 7B)**. We show quantification of basal respiration and basal glycolysis in all cell confluency groups. *Selenon*-KO myotubes exhibited significant decreases in OCR and ECAR in compared to WT when seeded at 20,000 cells/well **(Fig. 7C, D)**. Primary quadriceps myotubes from Selenon-KO mice *Selenon*-KO mice exhibited 78% reduction in OCR and 60% reduction in ECAR when compared to WT. The differentiated C2C12 cells with exon 3 *Selenon*-KO had 60% reduction in OCR and 61% reduction in ECAR. And lastly, C2C12 *Selenon*-KO exon 5 cells had 50% reduction in OCR and 61% reduction in ECAR. These data indicate that SelN-deficiency triggers metabolic deficiency at high myotube confluency. These observations further support mitochondrial involvement in *SELENON*-RM.

To test whether similar effects in metabolism might be detectable in vivo in our zebrafish models of *SELENON*-RM, we tested 1dpf embryos using the cell respirometer. We replicated the Seahorse protocol used for myotubes and tested homozygous embryos for each zebrafish line. In contrast to cultured and differentiated murine myotubes we did not detect measurable differences in respiration levels between WT and selenon-mutant embryos of any of the three independent lines **(Fig. 7E)**.

## DISCUSSION

A major challenge in the study of *SELENON*-RM has been that, despite the significant disease burden in patients afflicted with this condition, cellular and animal models do not exhibit clear phenotypes (Deniziak et al., 2007; Moghadaszadeh et al., 2013; Rederstorff et al., 2011). Although molecular and biochemical abnormalities related to SelN deficiency have been well documented, difficulties in demonstrating readily assayable clinical phenotypes have inhibited therapy development. Here we present three new zebrafish models for the study of *SELENON*-RM. Each line presents with different mutations in *selenon* exon 2 ensuring reproducibility of our findings. Two of our lines resulted in frameshift mutations: *selenon* ^cl502^ and *selenon*^cl503^ (c.197-200del and c.199-203del correspondingly); while the third resulted in an in-frame loss of one amino acid: *selenon*^cl504^ (c.197-199del). We demonstrate that our frameshift zebrafish models exhibit complete absence of SelN while our in-frame line has reduced levels of the abnormal protein **(Fig. 1)**. This in-frame mutation resulted in replacement of valine by glycine in amino acid position 66 and deletion of leucine at the 67 position (p.G66VdelL67). Glycine is reported as an unreactive amino acid while valine has a hydrophobic side chain. To further understand the implications of this mutation, we performed 3D modeling using the SWISS model algorithm. Results show changes in tertiary structure in SelN α-helix domain **(Supplemental Fig. 1)**. This suggests that this mutation in SelN results in reduced protein stability that may result in degradation by the cell.

Quantitative RT-PCR analysis of *selenon* mRNA levels at 24hpf and 6dpf in our lines revealed highest levels at 24hpf, decreasing by 6dpf in WT fry, suggesting an important role of *selenon* expression during early embryonic development. Frameshift-mutant fish had lower levels of steady state selenon mRNA at both 24hpf and 6dpf, perhaps reflecting the consequences of nonsense-mediated decay. However, our in-frame mutant fish showed increased expression at 6dpf, when compared to WT. This is consistent with a previous study by Maiti *et al.,*2009, where they measured SelN protein and *SELENON* transcript levels in biopsies from SELENON-RM patients. They found that when protein levels where null, transcript level was reduced compared to control (Maiti et al., 2009). However, when SelN protein levels were reduced, transcripts were elevated in patient samples when compared to controls, possibly indicting a potential compensatory response.

Measurement of spontaneous contractions (coiling) in zebrafish is a sensitive functional assay to measure embryonic muscle function (de Oliveira et al., 2021; Roussel et al., 2021). These contractions are mediated by slow-twitch oxidative fibers (Naganawa and Hirata, 2011) rich in mitochondrial content (McDermott and Bonen, 1992; Schiaffino and Reggiani, 2011). All three *selenon* mutant lines exhibited significantly decreased mean durations of contraction when compared to WT **(Fig 2)**. suggesting an impaired ability to maintain contractions in SelN deficient embryos. Additionally, the percent of activity was significantly reduced in two *selenon* mutants and showed a trend towards decrease in a third line (*selenon*^cl502^). This may indicate that because of the reduction in contraction duration, total activity declined. Finally, *selenon*^cl502/cl502^ embryos did not show differences in contraction count, while *selenon*^cl503/cl503^ did show a significant decrease, and *selenon*^cl504/cl504^ *in*-frame mutant showed increased count. Possibly SelN deficient embryos may be able to compensate for reduced contractile duration by increasing their count.

To test changes in swimming activity during early larval stages, we developed a light-and-dark cycle, and vibration protocol **(Fig.3)**. As expected, we found that all fish increased their activity in the dark and slowed down during light periods. We did not recognize any changes in patterns of behavior between *selenon* mutants and WT. However, we detected a significant decrease in total swimming distance in frameshift mutants *selenon*^cl502/502^ and *selenon*^cl503/503^ and a trend towards decreased distance in in-frame mutant *selenon*^cl504/cl504^ fry. Ultrastructural analysis of our zebrafish skeletal muscle did not identify any major changes to the myofibrillar ultrastructure, however we found differences in mitochondrial area sizes. We report increased area in both *selenon*^cl502/cl502^ and *selenon*^cl503/cl503^ and decreased mitochondrial size in *selenon*^cl504/cl504^ fish at 6dpf when compared to WT. These changes in skeletal muscle mitochondria along with changes in swim activity support the idea that SelN plays a role in skeletal muscle mitochondrial function during zebrafish early growth.

Given the increased evidence that SelN may play an important role in zebrafish embryos at 24hpf, we performed a bioinformatic analysis using previously published single cell RNAseq atlas (Wagner et al., 2018). We identified cell clusters that showed incremental expression of *selenon* gene from 6 to 24hpf, peaking at 24hpf. These clusters belong to the notochord and presomitic mesoderm (PSM) cells, precursors of the vertebral column and somites respectively (**Fig. 4A**). This, along with our previous data in 24hpf embryos, point at an important role for SelN in axial muscle development.

*SELENON*-RM patients exhibit rigidity of the spine and scoliosis but do not exhibit signs of neurogenic disease (Ferreiro et al., 2002; Moghadaszadeh et al., 2001; Saini et al., 2018; Scoto et al., 2011; Tajsharghi et al., 2005), so we decided to focus our attention on the presomitic mesoderm cells. Our analysis identified genes co-expressed with *selenon* from 6 to 24hpf. Gene ontology and pathway analysis suggests a strong correlation between *selenon* expression and glutathione redox reactions, and apelin adipocyte signaling pathways (**Fig. 4B-C**). However, the apelin adipocyte pathway contained the same participating genes as in glutathione redox pathways: *gpx7, gpx8, gpx4a,* and *gpx19*, likely accounting for that association. This is consistent with previous data implicating a role of SelN in redox reactions through the glutathione pathway (Arbogast et al., 2009; Chernorudskiy et al., 2020; Zito and Ferreiro, 2021).

To extend these observations we generated shRNA-induced *Selenon* knock down C2C12 cell lines (*Selenon*-KD) and quantified glutathione (GSH) and redox ratios (GSH_free_/GSSG) in control versus *Selenon*-KD cells, demonstrating a decrease in ratio proportional to their decrease in *selenon* expression **(Fig. 5A)**. Decreased levels of GSH_free_/GSSG can be interpreted as increasing concentration of oxidized glutathione when compared to the free available reduced glutathione. This happens in response to increased oxidative stress in the cells. These data may indicate a direct correlation between *selenon* expression and glutathione homeostasis in response to increased oxidative stress in myoblasts.

Additionally, we tested glutathione homeostasis in our *SELENON*-RM zebrafish models at 6dpf **(Fig. 5B)**. In contrast with C2C12 cells, we can normalize our data by number of fish, which allows us to report oxidized GSH (GSSG) and total glutathione levels (GSH_total_) per fish, as well as their GSH_free_/GSSG ratio. Results showed that fish with frameshift mutations in the *selenon* gene showed a significant decrease in GSSG. However, the in-frame *selenon* mutation line showed a significant increase in GSSG. Next, we showed that GSH_total_ was significantly decreased and showed a trend towards decrease in frameshift mutations correspondingly. The in-frame mutation model, however, showed a significant increase in total glutathione. Lastly, we showed that the redox GSH ratios in the three lines were not significantly different when compared to their WT lines. However, they showed a trend towards increased GSH_free_/GSSG in *selenon*^cl502^ and *selenon*^cl503^, and a trend towards decreased ratio in *selenon*^cl504^. These data may indicate that SelN deficient zebrafish have a heightened ability to adapt their GSH redox ratio when compared to isolated myoblasts. Lastly, we observe that *selenon*^cl502^ fish line presents the largest differences in GSSG and GSH_total_ when compared to *selenon*^cl503^. This corresponds to less reduction in spontaneous coiling and swim activity. This correlation may indicate that stronger GSH redox adaptations result in a milder phenotype.

To test whether oxidative stress is elevated in SelN deficient myoblasts, we measured reactive oxygen species (ROS) using 2, 7 -Dichlorodihydrofluorescein (DCFH) dye(Chen et al., 2010) **(Fig. 6A-B)**. We found that when *selenon* gene was significantly knocked down, DCFH fluorescence intensity went up when compared to control cells. We then wondered if this increased level in ROS rendered protein carbonylation, which has been shown to lead to contractile dysfunction in skeletal muscle and cell death **(Fig. 6C-D)** (Baumann et al., 2016). We also tested protein nitrosylation to determine whether NO damage was also present in SelN deficient cells. Our results show that protein carbonylation and nitrosylation were elevated in our *selenon-*KD cell lines. This is consistent with a previous study showing increased protein carbonylation in *SELENON*-RM patient myoblasts (Arbogast et al., 2009), supporting the concept that SelN plays a direct role in oxidative stress response in myoblasts.

The presence of minicores in muscle biopsies and impaired metabolic presentation in patients, suggest a direct relationship between SelN and mitochondrial function. However, mitochondrial abnormalities are a common secondary feature of muscle diseases. Here we demonstrate a direct relationship between mitochondrial dysfunction and SelN deficiency in muscle cells. We used CRISPR-Cas9 technology to generate two new C2C12 myoblast *Selenon* null lines. We then used a Seahorse cell respirometer to measure metabolic function in these cells after differentiation by measuring cellular respiration and glycolysis. We tested them in increasing concentrations of cells and found that at high confluency SelN-deficient myotubes show impaired metabolic function **(Fig. 7B)**. This may indicate that SelN’s role becomes more important during increased cellular stress. To prove that these metabolic changes were consistent, we tested our two *Selenon*-null C2C12 lines and primary cells isolated from *Selenon-*null mouse model quadriceps. We found that all lines showed metabolic impairment in both cellular respiration and glycolysis at high cell concentrations **(Fig. 7C)**. Being that we have previously shown that SelN may have a direct impact in oxidative stress, this data suggests that SelN-deficient oxidative stress leads directly to impaired mitochondrial function followed by changes in glycolysis. Interestingly, we failed to detect similar metabolic changes in 24hpf zebrafish embryos. This may indicate that metabolism is not impacted during somitogenesis perhaps due to the preponderance of fast oxidative myofibers in zebrafish and their reliance on glycolysis as primary source of energy(Jackson and Ingham, 2013; Miyazawa et al., 2022).

The present data provide new insights into the role of SelN in early embryonic development and provide the basis for new potential assays for drug and other therapeutic development. Further studies of mitochondrial structure and function in SelN deficient cells may provide a cellular morphological phenotype that might be of use in a cell painting type assay for drug screening (Cimini et al., 2023). Assays of spontaneous coiling and motor activity in zebrafish embryos and fry may have further utility for moderate throughput drug screens and in vivo proof of concept for other developing therapeutic modalities.

## METHODS

### Statistical analysis

Statistical calculations were performed using GraphPad Prism 10 software. Student’s *t* test was used to compare means between two groups of data and one-way ANOVA was used to compare three or more groups of data. ANOVA calculations were followed up by Tukey’s multiple comparison test to determine differences between samples. Means of experimental groups were considered significant when *p* value was < 0.05.

### Generation of *SELENON*-RM zebrafish models

To generate zebrafish lines with mutations in the *selenon* gene, we induced mutagenesis using CRISPR-Cas9 system Gene Art Platinum Cas9 Nuclease (ThermoFisher). This was achieved by performing mRNA microinjections into one-cell AB zebrafish embryos targeting exon 2. The guide RNA (gRNA) used to target this exon was 5’-AUU GUA GGA GGC AGG ACU GA-3’. This resulted in the generation of mosaic mutations in exon 2 of the *selenon* gene. These eggs were then reared and bred to AB wild type (WT) to generate heterozygous carriers. After Sanger sequencing and selecting mutations of interest fish were outbred to AB WT. Finally, each heterozygous mutation was intercrossed to generate experimental fish: homozygous knock outs (KO), heterozygotes (HET), and WT in ratios of 1:2:1.

### Zebrafish Real Time qPCR

We quantified *selenon* transcripts in zebrafish using PrimePCR multiplex real time quantitative PCR (RT-qPCR) system by Bio-Rad. We extracted and purified total RNA from cells and zebrafish using Trizol LS reagent (Ambion) following protocol by Peterson et al., 2009 (Peterson and Freeman, 2009). Reverse transcription and qPCR assay was performed using Reliance One-step multiplex RT-qPCR Master Mix (BioRad). We included two housekeeping genes determined by Reference Gene H384 panel by Bio-Rad: *rps18* and *ppia*. Analysis was performed using CFX Maestro Software by Bio-Rad.

### Zebrafish Western Blotting

For protein quantification of SelN in zebrafish embryos, we created new antibodies against the C-terminal domain of SelN for both zebrafish and mouse isoforms (Biomatik). Cells and 2dpf deyolked and dechorionated zebrafish (according to (Link et al., 2006)) were extracted for total protein using RIPA lysis buffer (Thermofisher) with protease inhibitors (Thermofisher). Using NuPAGE-Tris Acetate western blot system by Invitrogen we separated our proteins by size and transferred them onto a PDVF blot. Ponceau stain (Sigma Aldrich) was used to image total protein transferred. Finally, we immunoblotted against SelN at a 1:500 dilution and rhodamine labeled anti-tubulin antibody (Bio-Rad) at 1:5,000 concentration. Blots were imaged using a ChemiDoc MP Imaging System by Bio-Rad.

### Zebrafish spontaneous coiling assay

To measure spontaneous coiling, we set up zebrafish breeding pairs overnight with dividers and began mating the following day. Fish were harvested in windows of 30 min and time was annotated to maintain consistency. 24 hours later, embryos were placed under a light microscope and recorded for 3 min. Cycles of ∼15 embryos were recorded alternating mutant and WT fish for one hour. To quickly fit ∼15 embryos under the microscope field of view we 3D printed a 1.4 mm wide grid. Videos were analyzed using DanioScope software.

### Zebrafish hatching assay

To measure changes in hatching we set up HET by HET zebrafish breeding pairs overnight with dividers and began mating them the following day at the same time. One hour later we harvested the eggs and allowed them to grow overnight. On day 1 post-fertilization eggs were transferred to three 96-well plates for observation. Hatching counts were performed 4 times a day until all eggs had hatched. On 4dpf fish were euthanized and genotyped.

### Zebrafish swim activity assay

6dpf zebrafish larvae were placed in 48-well plates and tested for swim activity using an activity monitor by Zantiks. The protocol used during activity recording included: 10 min acclimation, 20 seconds vibration, 5 min light, 5 min dark, 5min light, 5min dark, and then repeat the same steps. Data was analyzed using in-house generated MatLab script (see **Supplemental File 3**).

### Analysis of embryonic zebrafish single cell RNAseq Atlas

Single-cell RNAseq (scRNAseq) data from Wagner *et al*. 2017 were downloaded. We extracted gene expression data using the Python package scanpy v1.9.1. In total, 30677 genes with their averaged expression data across six samples in the scRNAseq data were extracted. 903 genes were removed during QC step for having missing data and/or zero variance. Expression matrix was loaded on R version 4.0.2 for analysis. Adjacency matrix for the remaining 29774 genes was generated using the Weighted Gene Correlation Network Analysis (WGCNA) package (Langfelder and Horvath 2008). Soft thresholding from the scale-free topology model was used to reduce ‘noise’ of correlations in the adjacency matrix. A Topology overlap matrix (TOM) was generated from the adjacency matrix and converted to a dissimilarity matrix by subtracting the TOM matrix from 1. Samples were clustered using the ‘hclust average’ method. 136 gene modules were identified using the dynamic tree cut method on the generated clusters using the WGCNA R package with a minimum of 30 genes per module. To distinguish between these modules were labeled as colors using the labels2colors function. 20 core merged modules were generated by merging modules with a similar matrix (distance threshold < 0.25). The genes in each module demonstrate a similar expression profile and co-express. Pearson correlation was generated for each module against clinical traits, i.e., time point. Significance of the correlation of each gene against the time point was computed. Two core modules, ‘antiquewhite1’ (4516 genes) and ‘firebrick2’ (240 genes) having target selenon gene with significant p-values < 0.05 were identified. 3339 genes with positive association (weighted threshold > 0.02) to S*elenon* (sepn1) from ‘antiquewhite1’ and 168 genes from ‘firebrick2’ modules were exported using the exportNetworkToCytoscape function and were used to generate interaction networks on CytoScape v3.8.2 software. KEGG pathway and GO term enrichment analysis of significant modules was performed using clusterProfiler package (v3.16.0) (Yu et al., 2012). Enriched pathways were identified as terms with p.adj < 0.05.

### Cell culture

C2C12 mouse myoblasts (American Type Culture Collection, Manassas, VA) were maintained in growth medium (DMEM supplemented with 20% Fetal Bovine Serum, Certified; GIBCO, (Invitrogen, Carlsbad, CA, USA) in 10% CO2. For differentiation, cells were allowed to grow to 90-100% confluence and switched to differentiating medium (DMEM containing 2% Horse Serum; GIBCO, Invitrogen, Carlsbad, CA, USA) that was subsequently changed every 24 h.

### C2C12 *Selenon* knock down

RNAi-Ready pSIREN-RetroQ vector (Clontech, Inc. Mountain View, CA, USA) was used to prepare shRNA constructs targeting *Selenon* gene (KD1: TTCAAACCCATTGCGGAGA, KD2: GCAAACCATGAATTGGAAAGT, and KD3: CATGATTGACAGCCGCCTG). These constructs were transfected into the 293 Ecopack packaging cell line (Clontech) using Lipofection 2000 transfection reagent (Invitrogen, Carlsbad, CA, USA) to generate retrovirus particles. 48 hours after transfection, the viral supernatant was harvested and used for transduction of C2C12 cells. The virus was removed 6 hours after the infection and fresh growth medium containing puromycin was added to C2C12 cells allowing the selection of infected cells. For each shRNA target 3 replicate plates of C2C12 cells were transduced and were subsequently processed as independent cultures.

### Cellular RNA extraction and qRT-PCR

Total RNA was extracted and purified from C2C12 cells using RNeasy kit (Qiagen, Valencia, CA) according to the manufacturer’s instructions. RNA was treated with DNase I Amplification Grade (Invitrogen, Carlsbad, CA, USA). cDNA synthesis preparation was performed using the SuperScript First-Strand synthesis system for RT-PCR (Invitrogen, Carlsbad, CA, USA). Real-time quantitative RT-PCR was carried out using the TaqMan probe-based chemistry (Applied Biosystems, Foster City, CA, USA) on an ABI Prism 7300 Real Time PCR System.

### Cellular Western Blotting

Cell cultures in 100 mm dishes were washed twice with PBS and scraped into 300 μl of sample buffer. Lysates were denatured for 5 min at 95°C, sonicated with Branson Sonifier 250 to shear DNA and centrifuged for 10 min at 16,000 rcf. Soluble proteins were quantified by Lowry’s method using the D_C_ Protein Assay Kit (Biorad, Hercules, CA, USA). Twenty micrograms of whole cell extracts were resolved on 4−12% SDS-polyacrylamide gels and transferred onto Ready gel blotting sandwiches immune-blot PVDF membranes (Biorad, Hercules, CA, USA) using a XCell II™ Blot Module (Invitrogen, Carlsbad, CA, USA). Membranes were then probed with crude SelN antibody 7637 raised in rabbit (1:500). After incubation with HRP-conjugated AffiniPure Goat Anti-Rabbit IgG (1:10,000), reactive proteins were visualized with Supersignal West Pico Chemiluminescent substrate (Fisher Scientific Pittsburgh, PA, USA).

### Glutathione assay in cells and zebrafish

Zebrafish and cells were tested for oxidized and reduced forms of glutathione (GSH) molecule using glutathione colorimetric detection kit by Thermo-Fisher. Five 6dpf zebrafish per sample were humanely euthanized and homogenized using 100ul of 100mM phosphate buffer, pH7. Procedure for both cell and zebrafish was continued as indicated by the manufacturer. Colorimetric reaction was measured using a plate reader every minute for 20 min.

### ROS quantification in C2C12

Intracellular ROS was measured using CM-H_2_DCFDA (Invitrogen, Carlsbad, CA, USA). Briefly 3 samples per group of cells were detached by trypsin and washed in PBS and spun down. The cell pellets were then resuspended in 100 μl of PBS containing 5 μM CM-H_2_DCFDA and incubated at 37°C for 15 minutes. Cells were then washed with PBS and resuspended in 0.5ml of PBS. 10,000 cells per sample were analyzed for fluorescence using Becton Dickinson FACS-Vantage SE flow cytometer. The data were analyzed using Flowjo software (Tree Star, Inc., OR, USA).

### Carbonylation and nitrosylation assay

Total carbonylated proteins were quantified using OxyBlot Protein Oxidation Detection Kit (Millipore, Billerica, MA) following manufacturer’s instructions using 15 μg of total protein detected with rabbit anti-DNP antibodies (Invitrogen) at 1:1,000 and HRP-conjugated AffiniPure Goat Anti-Rabbit IgG (1:10,000). Total S-nitrosylated proteins were measured using an S-nitrosylated Protein Detection Assay Kit (Cayman Chemical Company, Ann Arbor, MI, USA). Both carbonylated and S-nitrosylated proteins were detected using Supersignal West Pico Chemiluminescent substrate (Fisher Scientific) and quantified by scanning the gels on a ChemiDoc XRS gel documentation system and analyzed using Quantity One software v. 4.5.2 (Bio-Rad). The top of each gel was defined as Relative front (Rf) = 0 and the bottom as Rf = 1. Repeated intensity measurements were made down the center of each lane at intervals of Rf = 0.0043.

### Generation of *Selenon* knock out C2C12 cell lines

Using CRISPR-Cas9 system we stablished two new *Selenon*-null C2C12 myoblast lines. These lines were generated independently. One resulted in mutations in exon 5 of the *Selenon* gene, and the second one in exon 3. This was achieved by using lipofectamine and transfecting with LentiCRISPRv2 plasmids. The following sequences were used to clone gRNAs coding regions in the plasmids for *selenon* exon 3 or exon 5 gRNA’s: Ex3-F 5’-caccgTATGGTAAGAGTCTCCTCGC-3’, Ex3-R 5’-aaacGCGAGGAGACTCTTACCATAc-3’, Ex5-F 5’-caccgGGCAAAGCGGGTCTTCACGA-3’, and MouseSepnEx5-gRNA-R 5’-aaacTCGTGAAGACCCGCTTTGCCc-3’. Puromycin antibiotic was used to select for transfected cells and single cells were isolated and grown individually. New single colonies were expanded and sequenced to identify null mutations.

### Primary Myoblast Isolation

To isolate primary myoblasts from our *SELENON*-RM mouse model, we euthanized and dissected the quadricep muscles from 3-month-old mice. We then followed myoblast isolation protocol by Shahini et al., 2018 (Shahini et al., 2018). Myoblasts were frozen in 10% DMSO growth media and utilized up to 7 passages.

### Seahorse (Agilent) metabolic function in myotubes and zebrafish embryos

To test metabolic function in C2C12 cells, primary cell lines, and 24hpf zebrafish embryos we used the Seahorse XFe96 Analyzer by Agilent following manufacturer instructions. For zebrafish experiments we placed a single dechorionated embryo in XFe96 Spheroid Microplate per well in Danio water and measured real time metabolic activity using the following protocol: Port A injected a final concentration of 12.5uM oligomycin, port B injected 2uM FCCP and 1mM sodium pyruvate, port C injected 10uM rotenone and antimycin-A, and port D injected 5mM 2-deoxyglucose. Timing between measurements was: 2 min mixing, 1 min wait, and 2 min measure, repeated 5 times for baseline and after each port injection.

C2C12 cells and primary myoblasts, cells were seeded onto XFe96 Cell Culture plate precoated with 0.09mg/ml Matrigel (Corning, NY). The following day cells were differentiated using 2% horse serum media for C2C12 and according to (Shahini et al., 2018) instructions for primary cells, for 5 days. Myotubes were then tested in the Seahorse XFe96 Analyzer using the following protocol: Port A injected a final concentration of 1.5uM oligomycin, port B injected 1.5uM FCCP and 1mM sodium pyruvate, port C injected 2.5uM rotenone and 1.5uM antimycin-A, and port D injected 5mM 2-deoxyglucose. Timing between measurements was: 2 min mixing, 1 min wait, and 2 min measure, repeated 3 times except for measurements after port B, which were repeated 5 times. Immediately upon assay completion, leftover media was carefully removed, and cells were lysed for creatine kinase assay and protein concentration for normalization. Colorimetric creatine kinase activity assay (Abcam) was performed after each Seahorse assay followed by BCA protein assay (Fisher Scientific).

### Ultrastructure imaging of zebrafish using Transmission Electron Microscopy

6dpf zebrafish larvae were euthanized and placed in glutaraldehyde fixative diluted in fish water at a 1:1 ratio. After 2 days, fish were sectioned and prepared for transmission electron microscopy (TEM) imaging using JEOL 1200EX-80kV microscope. Slow twitch fibers were identified as adjacent to the skin and myofibrils were imaged at 2500X. At least 13 images were recorded for each fish and mitochondria areas were quantified using Image J software.

## Acknowledgements

Thanks to Elizabeth Griffin for outstanding technical assistance during a summer internship. The authors would also like to acknowledge helpful support and advice from Louise Trakimas of the Electron Microscope Core at Harvard Medical School, as well as Shanelle Kohler from Zantiks Ltd. Electron Microscopy Imaging, consultation and /or services were performed in the HMS Electron Microscopy Facility, and we would also like to acknowledge the Zebrafish Facility at Boston Children’s Hospital for their continuous care of our fish.

