## Supplemental Figures for "Zebrafish and cellular models of *SELENON*-Related Myopathy exhibit novel embryonic and metabolic phenotypes"

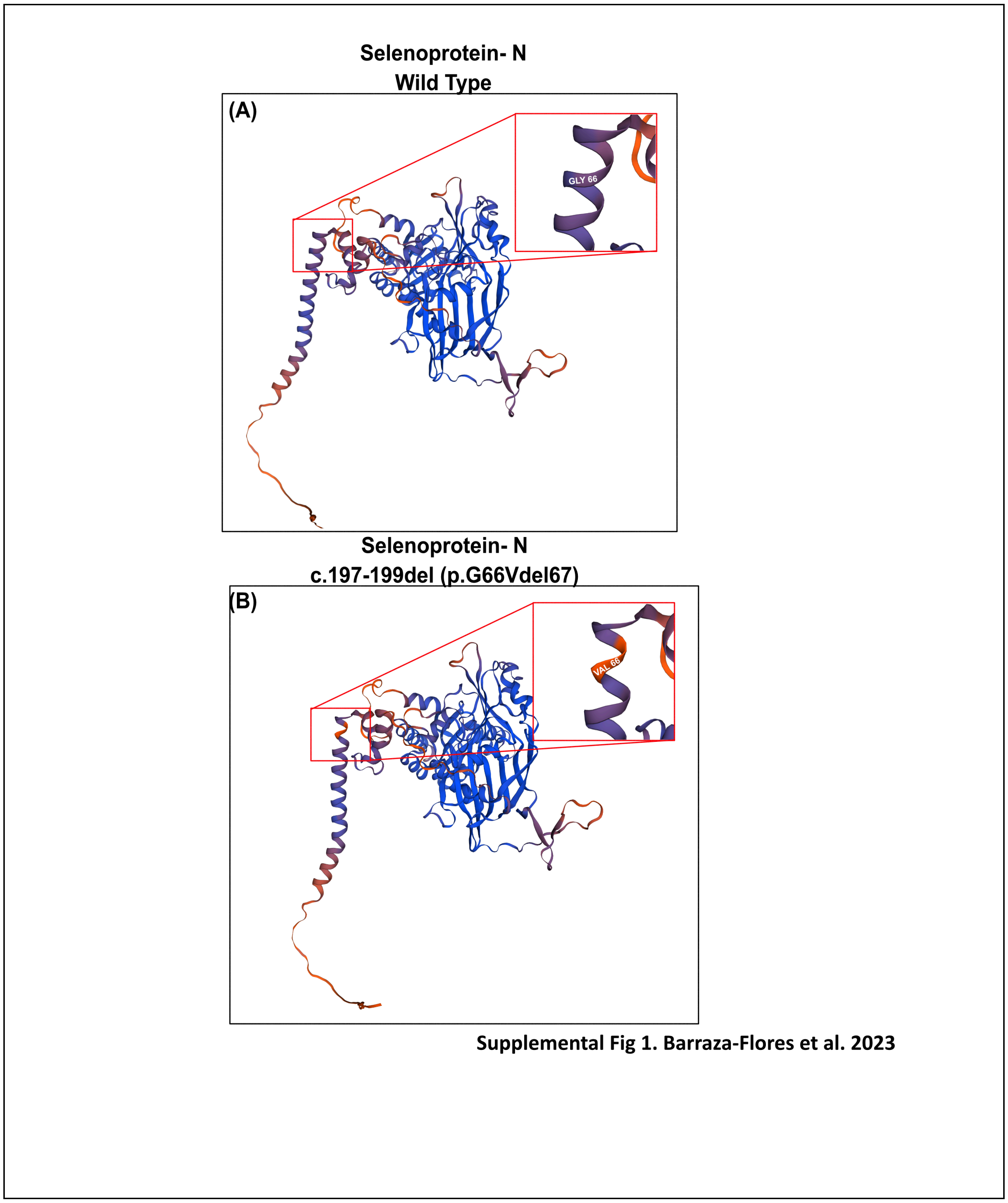


**Supplemental figure 1. *In silico* 3D modeling of wild type and mutant Selenoprotein-N.**

SWISS-MODEL web-based tertiary structure models of **(A)** wild type and **(B)** mutant c.197-199del Selenoprotein-N with amino acid sequence mutation p.G66VdelL67. Color scheme represents protein confidence score with orange being very low (score<0.5) and blue very high (score>0.9) . Red squares amplify $\alpha$-helix subunits at the site of mutation.

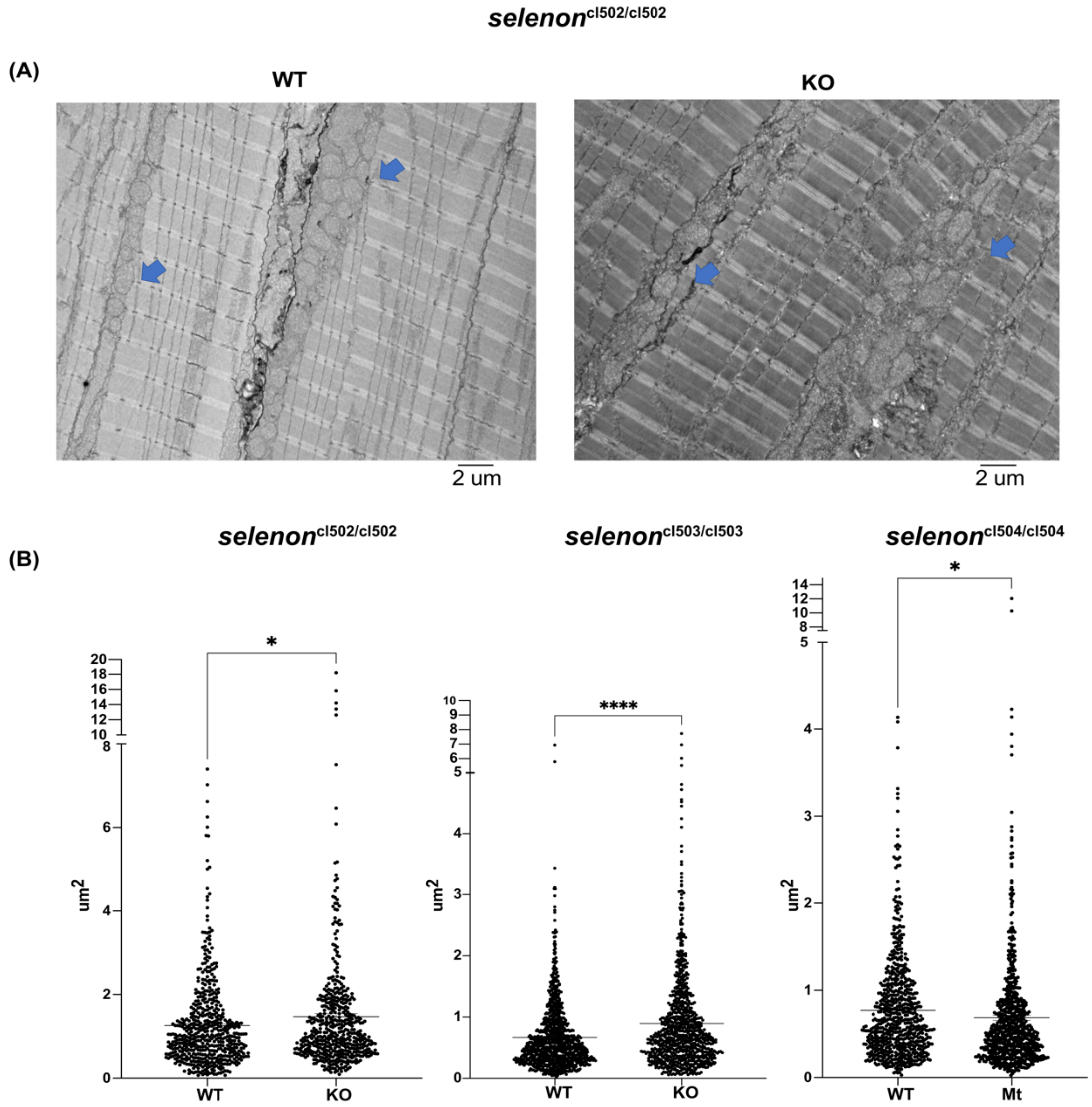


Supplemental Fig.2 Barraza-Flores et al. 2023

**Supplemental figure 2. *Selenon*-deficient zebrafish present with changes in mitochondrial morphology when compared to WT. (A)** Electron microscopy images of longitudinal slow twitch muscle fibers' ultrastructure from WT and *selenon* deficient zebrafish at 6dpf (scale bar 2 μm). Mitochondria are shown by blue arrows. **(B)** Results of quantified mitochondrial area shows significant increases in homozygous *selenon*-null zebrafish, and a slight decrease in the selenon$^{cl504/cl504}$ mutants when compared to WT (N=2). “*” = $p<0.05$, “****” = $p<0.0001$.

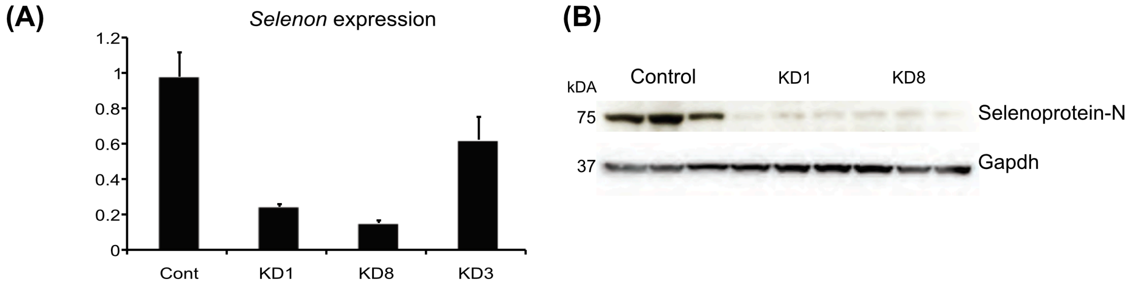


Supplemental Fig. 3. Barraza-Flores et al. 2023

**Supplemental figure 3. Selenon expression in knocked down C2C12 cell lines. (A)** *Selenon* transcript levels demonstrates significant decreased expression in KD1 and KD8 lines. **(B)** Western blot demonstrates decreased levels of Selenoprotein-N in KD1 and KD8 lines.

**Supplemental Table 1.** 'Oxidoreductase activity acting on peroxide as an acceptor' pathway gene list from gene ontology analysis of 'Antiquewhite1' module. Genes are organized by weight of correlation to selenon expression pattern.

| Gene ID | Weight | Gene Symbol | Description |
|---|---|---|---|
| 552981 | 0.4499004 | *gpx7* | Glutathione peroxidase7 |
| 570477 | 0.44628944 | *prdx4* | Peroxiredoxin 4 |
| 405772 | 0.42508608 | *hbbe2* | Hemoglobin beta embryonic-2 |
| 393191 | 0.42191258 | *gpx8* | Glutathione peroxidase 8 |
| 554105 | 0.41837226 | *prdx5* | Peroxiredoxin 5 |
| 352928 | 0.4083269 | *gpx4a* | Glutathione peroxidase 4a |
| 30507 | 0.40275171 | *hbaa1* | Hemoglobin alpha adult 1 |
| 30596 | 0.39145277 | *hbbe3* | Hemoglobin beta embryonic 3 |
| 30601 | 0.39141568 | *hbae3* | Hemoglobin alpha embryonic 3 |
| 337376 | 0.37776591 | *alox5ap* | Arachidonate 5-lipoxygenase-activating protein |
| 791455 | 0.37737733 | *prdx2* | Peroxiredoxin 2 |
| 352926 | 0.34475496 | *gpx1a* | Glutathione peroxidase 1a |
| 100150283 | 0.29270673 | *loxhd1b* | Lipoxygenase-homology PLAT domains 1b |
| 436833 | 0.18566916 | *gstk1* | Glutathione S-transferase kappa 1 |
| 564819 | 0.03973992 | *loxhd1a* | Lipoxygenase-homology PLAT domains 1a |
